# A multi-phase flux balance model reveals flexibility of guard cell central carbon metabolism

**DOI:** 10.1101/2020.01.13.905372

**Authors:** Joshua XL Tan, C. Y. Maurice Cheung

## Abstract

Experimental research in guard cell metabolism has revealed the roles of the accumulation of various metabolites in guard cell function, but a comprehensive understanding of their metabolism over the diel cycle is still incomplete, given the limitations of current experimental methods. In this study, we constructed a four-phase flux balance model of guard cell metabolism to investigate the changes in guard cell metabolism over the diel cycle, including the day and night and stomata opening and closing. Our model demonstrated the metabolic flexibility in guard cells, showing that multiple metabolic processes can contribute to the synthesis and metabolism of malate and sucrose as osmolytes during stomatal opening and closing. We showed that guard cells can adapt to varying light availability and sucrose uptake from the apoplast during the day by operating in a mixotrophic mode with a switch between sucrose synthesis via the Calvin-Benson cycle and sucrose degradation via the oxidative pentose phosphate pathway. During stomatal opening, our model predicted an alternative flux mode of the Calvin-Benson cycle with all dephosphorylating steps diverted to diphosphate—fructose-6-phosphate 1-phosphotransferase to produce PP_i_, which is used to pump protons across the tonoplast for the accumulation of osmolytes. An analysis of the energetics of the use of different osmolytes in guard cells showed that malate and Cl^-^ are similarly efficient as the counterion of K^+^ during stomatal opening.

**Significance statement:** This work presents the first four-phase metabolic model for predicting guard cell metabolism over the diel cycle, which predicted an alternative flux model of the Calvin-Benson cycle that maximises the production of PP_i_ during stomatal opening. While multiple metabolic processes were shown be important in synthesising and metabolising osmolytes in guard cells of different experimental systems, our model demonstrated that these processes can operate simultaneously and at different rates depending on conditions.

## Introduction

The guard cell’s role of regulating gas and water exchange between a plant and the atmosphere makes understanding guard cell function crucial for a holistic understanding of the whole plant (Lawson et al., 2014; Azoulay-Shemer et al., 2016). Moreover, the recent recognition of the stomata’s impact on global water and carbon cycles (Azoulay-Shemer et al., 2016; Jezek and Blatt, 2017), their centrality to the projected water availability and crop production crisis in 20-30 years (Jezek and Blatt, 2017), and the resulting urgency to engineer water-efficient crops through altering stomatal function (Laporte et al., 2002; Lawson et al., 2014) makes a complete understanding of guard cell metabolism increasingly urgent.

We now have a broad understanding of how guard cells perform their physical function (outlined in numerous studies summarized by Lawson et al., (2014) and Santelia and Lawson (2016)). During stomatal opening, the uptake or synthesis of osmotically active metabolites (hereafter “osmolytes”) increases osmotic pressure within guard cells, causing the cell to become turgid and the stomatal pore to open (Outlaw, 2003). During stomatal closing, osmolytes are exported through efflux or degraded through metabolism, and guard cells lose turgor, thereby closing the stomatal pore (Outlaw, 2003). Still, certain aspects of guard cell metabolism remain open questions. Specifically, questions about the interaction between metabolism and osmolyte synthesis/transport remain an open area of study. For instance, what is the role of carbohydrate metabolism in guard cell function (Santelia and Lawson, 2016) and what is the origin of guard cell sugars (Lawson et al., 2014)? What is the rationale behind the choice of counter-ion? To what extent do photosynthetic products contribute to osmolyte production or energy for membrane transport (Santelia and Lawson, 2016)?

These questions implicate metabolic processes that are dynamic, involving metabolic processes that interact dynamically with one another, and with membrane transport reactions (Lawson et al., 2014). Experimental observations have provided measurements of guard cell metabolite and osmolyte accumulation in response to environmental stimuli (Santelia and Lawson, 2016; Santelia and Lawson, 2016) and accumulation of specific metabolites over a given time interval (Horrer et al., 2016). While experimentation has revealed the presence of interactions between metabolic processes (Lawson et al., 2014), it is limited in its ability to demonstrate the extent of the interactions, and to revealing the dynamics of these relationships over time (Jezek and Blatt, 2017). Moreover, as Jezek and Blatt (2017) pointed out, experimental manipulation of guard cells affects the balance of cytosolic interactions, limiting our ability to gain a holistic understanding of metabolic processes. Clearly, we need a method, complementary to experimental studies, that allows us to place experimentally observed facts about guard cell metabolite accumulation in the context of the cell’s entire metabolism, and one that enables us to see how these relationships change over time.

The application of flux balance analysis (FBA) on genome-scale metabolic models has emerged as a viable tool for investigating the behaviour of multiple plant metabolic systems (Sweetlove and Ratcliffe, 2011). FBA can account for systems-level interactions between various metabolic pathways (Sweetlove and Ratcliffe, 2011), and can model temporal behaviour in the form of interactions between multiple steady-states, using the diel modelling approach developed by Cheung et al. (2014). Moreover, condition-specific scenarios can be explored using FBA by setting corresponding constraints (Sweetlove and Ratcliffe, 2011). Robaina-Estévez et al. (2017) demonstrated the viability of constraints-based modelling in generating a functional large-scale analysis of guard cell metabolism, using transcriptomics data to define constraints for the genome scale model. In their study, differential analysis between the predicted fluxes of modelled guard cells and mesophyll cell model revealed key roles for the tricarboxylic acid (TCA) cycle in malate synthesis, the Calvin-Benson cycle in sucrose synthesis, and chloroplasts for guard cell function and metabolism (Robaina-Estévez et al., 2017). Their study shed light on the possible ways central carbon metabolism can structure the physiological function of guard cells. Our study extended beyond the characterisation of guard cell metabolism during stomatal opening with the addition of simulating metabolism through the whole diel cycle. We constructed a four-phase model of guard cell metabolism to investigate the central carbon metabolism in guard cells over multiple phases of a diel cycle.

## Results and Discussion

### A four-phase modelling framework for guard cell metabolism

To model the metabolism of guard cells over a diel cycle, including the opening and closing of stoma, a four-phase modelling framework was developed as an extension of a diel modelling framework (Cheung et al., 2014) to represent four distinct phases of guard cell metabolism. A core plant metabolic model from Shameer et al. (2018), which was used for modelling C_3_ and CAM photosynthesis, was converted into a single-phase model before being duplicated four times to produce a four-phase metabolic model of guard cell (Appendix S1, S2). The four-phases, Open, Day, Close and Night, correspond to the opening of stoma during sunrise, daytime, the closing of stoma during sunset and night-time respectively (Figure 1). The “Open” and “Close” phases are assumed to be each one hour in length, and the “Day” and “Night” phases are each eleven hours in length. Thus the diel cycle of the four-phase model consists of a 1-h/11-h Open-Day light phase, followed by a 1-h/11-h Close-Night dark phase.

**Figure 1.**
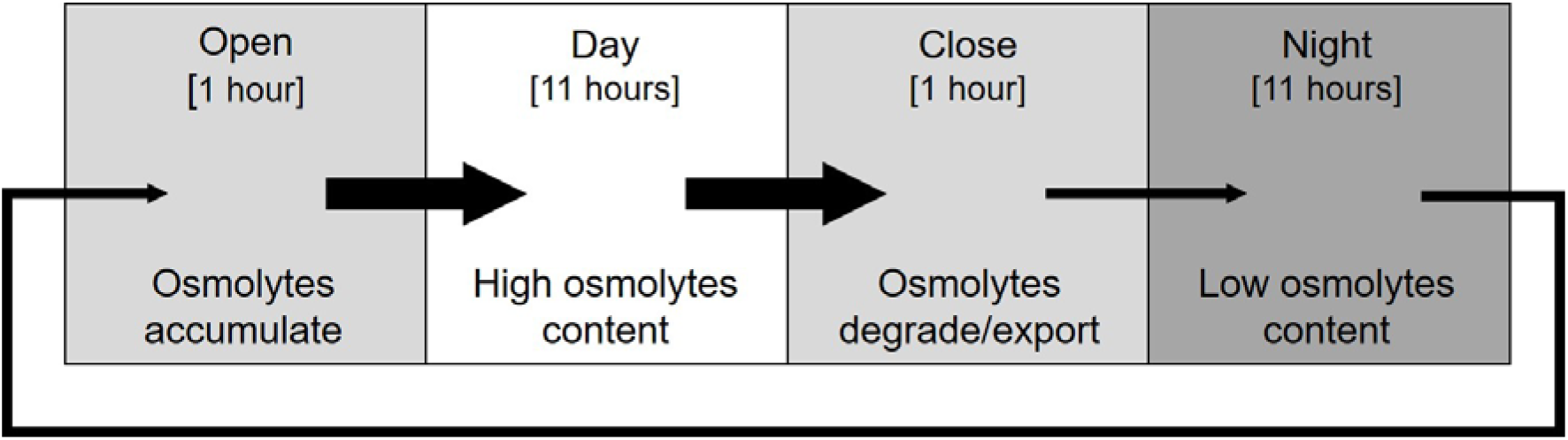
Four-phase model of guard cell metabolism. Each square represents a metabolic model of each phase. Black arrows represent the storage of metabolites from one phase to another phase. Thick arrows represent storage of metabolites including osmolytes; Thin arrows represent storage of metabolites excluding osmolytes.

In the four-phase modelling framework, the metabolism in each individual phase was assumed to be constant, i.e. in a steady-state. All four phases contain the same set of reactions except for phase-specific constraints applied to each phase. The interactions between two consecutive phases was modelled by reactions simulating the accumulation and degradation of storage metabolites (termed “storage reactions”), similar to the approach used in Cheung et al. (2014). However, whereas Cheung et al. (2014) utilised bidirectional storage reactions, the storage reactions in the present four-phase model are unidirectional. Storage reactions made the metabolites accumulated at the end of each phase available for consumption in the subsequent phase. Two main types of metabolites were selected as storages between the phases: osmolytes (K^+^, Cl^-^, malate, and sucrose) and carbon and energy reserves (starch and sucrose). Our modelling approach differs from the Robaina-Estevez et al. (2017) guard cell model in significant ways. Robaina-Estevez et al. (2017) used transcriptomics of guard cells and mesophyll cells to define the initial set of constraints on their modified AraCore model to model one particular phase during the day. Our selected constraints for our four-phase model were based on observed phase-specific metabolic behaviour (i.e. certain metabolites were known to accumulate during specific phases). This allowed us to model the metabolic changes and the pattern of metabolite and osmolyte accumulation over multiple phases in the diel cycle.

### Four-phase metabolic model successfully simulated known features of guard cell metabolism

The four-phase model was tested by setting a standard set of constraints to model guard cell metabolism, which we refer to as the “standard scenario”. The choice of constraints for the standard scenario was guided by three major considerations: i) the environment to which a guard cell is exposed to, ii) the opening and closing of stoma, and iii) other physiological functions of guard cell. The key constraints for guard cell metabolism are highlighted below. More detailed constraints are described in Material and Methods.

The first set of constraints took into consideration light availability and gaseous exchange. During the dark phases (Close and Night), photon influx was constrained to zero. During the light phases (Open and Day), the upper bounds of photon influx reactions was unconstrained, allowing photons to be used as energy sources for the guard cell. For gaseous exchange, the transport reactions for CO_2_ and O_2_ were set to be unbounded.

The second set of constraints took into consideration the change in osmotic pressure of the guard cells throughout the course of the day. To simulate the opening of the stomata during the Open and Day phases, we constrained the model to undergo a fixed change in osmotic pressure by setting a fixed sum of accumulation of osmolytes (0.9 mosmol g^-1^ DW based on Rechmann et al. (1990)), scaled by their osmotic coefficients (see Materials and Methods). To simulate the closing of the stomata during the Night and Close phases, we set the flux of osmolyte accumulation to zero between the day-to-close and close-to-night phases. Multiple studies provide experimental evidence for the accumulation of K^+^ in guard cells after the first few hours of light exposure (Tallman and Zeiger, 1988; Santelia and Lawson, 2016) with sucrose replacing K^+^ as the primary osmolyte in the latter part of the light phase (MacRobbie, 1987; Santelia and Lawson, 2016). To simulate the choice of osmolytes corresponding to specific times of day, we set K^+^ storage to zero during the Day phase and sucrose storage to zero during the Open phase. These constraints forced K^+^ to be exported after the Open phase and for sucrose to be accumulated during the Day phase. To maintain electroneutrality, the sum of osmolytes with electric charge (K^+^, Cl^-^, and malate) were constrained such that there was no accumulation of net charge. In the standard scenario, the use of Cl^-^ as osmolyte was disabled to simplify the interpretation of model predictions. The effect of the Cl^-^ constraint on model prediction was explored in section “Malate and Cl^-^ are similarly energetically efficient as counterion of K^+^”. Given the large uncertainty regarding the contribution of guard cell photosynthetic carbon reduction for osmolyte accumulation in stomatal opening, which ranges from 2% to 40% (Lawson, 2008), we assumed a value of 20% in the standard scenario. Based on this, we set sucrose import from the apoplast during the Day phase to be 80% of the osmolyte required for stomatal opening, i.e. 0.72 mmol sucrose g^-1^ DW, given the lack of knowledge of the contribution of external sucrose to guard cell metabolism. A sensitivity analysis of external sucrose import was performed and presented in the section “Changes in central carbon metabolism with external sucrose import into guard cells”.

The third set of constraints took into consideration the other physiological functions of guard cells. Guard cells do not produce biomass, so all biomass fluxes in the model were constrained to zero. Cellular maintenance was modelled similar to previous studies (Cheung et al., 2014; Shameer et al., 2018; see Materials and methods for detail). Given the lack of experimental information on guard cell maintenance costs, we varied the magnitude of cellular maintenance costs and selected the value of 2.87 mmol ATP g^-1^ DW hr^-1^ as the value for ATP maintenance cost for the standard scenario based on matching carbon fixation during the Day phase contributing to 20% of osmolyte accumulation (Lawson, 2008).

With the constraints for the standard scenario, we were able to obtain an optimal flux solution with osmolyte and starch accumulation patterns matching qualitatively with experimental observations in guard cells (Figure 2; Data S1). As expected from the constraints we set, the model simulated K^+^ and malate accumulation in the morning and sucrose accumulation in the afternoon. Despite not setting any constraints for starch accumulation, our model successfully predicted starch accumulation during the Day and Close phase and starch degradation during the Night and Open phase, which is consistent with experimental measurements of starch content in guard cells (Horrer et al., 2016). When the K^+^ and sucrose accumulation constraints were removed, the model predicted the use of K^+^and malate over sucrose as the preferred osmolytes over both Open and Day phases given the objective function of minimising photon influx (Data S1). The model required less photon if K^+^ and malate are allowed to accumulate in all phases, demonstrating that the use of K^+^ and malate as osmolytes is energetically more favourable than sucrose.

**Figure 2.**
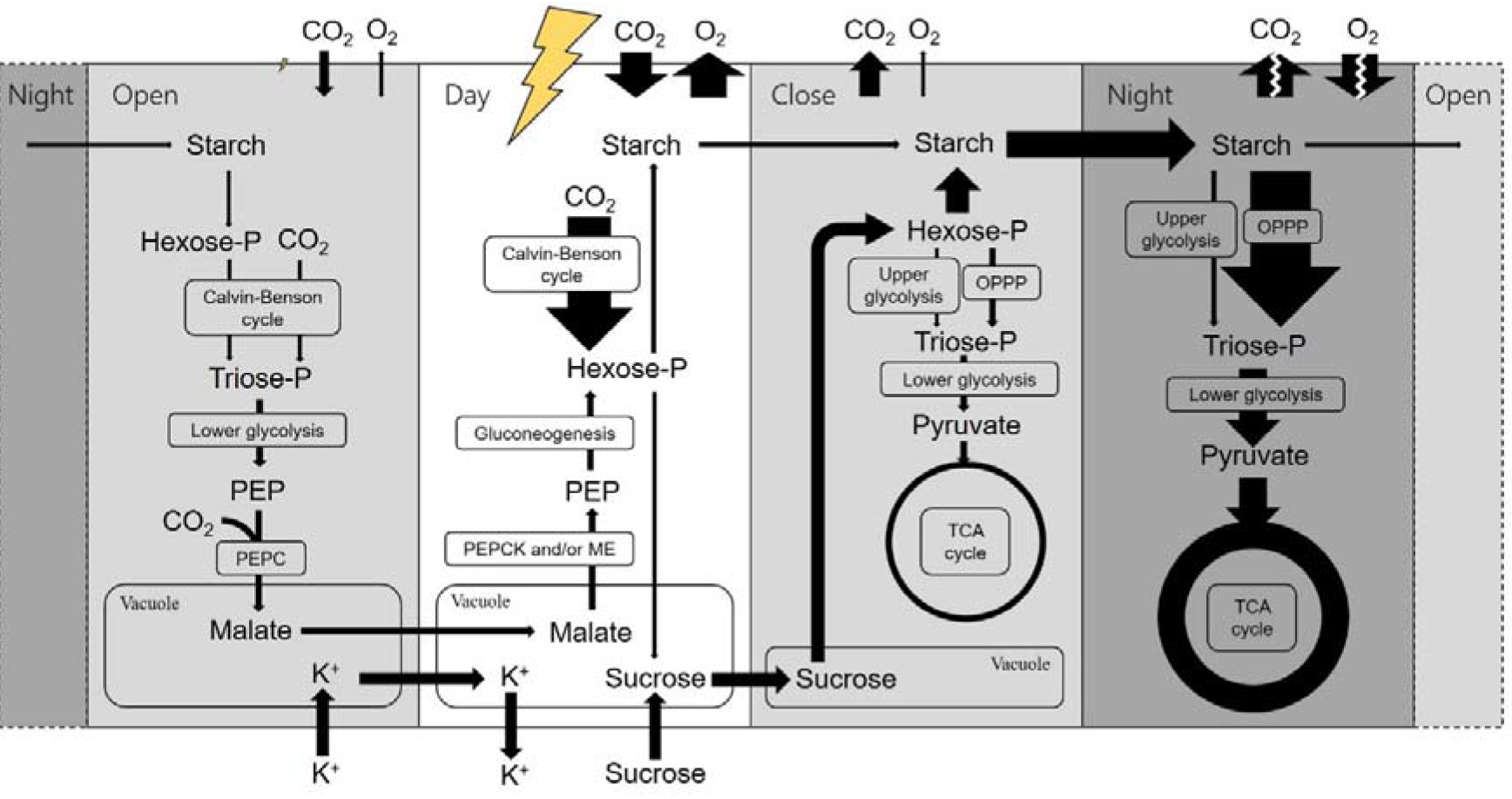
Flux map of osmolyte movements and carbon metabolism of guard cell under the standard scenario. Arrows connecting compounds within individual rounded rectangles denote reactions that occur within intracellular compartments or phase (according to labels). Arrows connecting compounds across phases denote storage compounds from the previous phase, made available for use in the next phase. Width of arrows reflects the relative magnitude of fluxes through the metabolic process scaled to flux per phase. Broken arrows represent fluxes too large to be shown in scale. Yellow lightning symbols represent photon influx per phase (not scaled relative to other arrows). The photorespiratory pathway and CO_2_ released from metabolic processes were omitted for clarity. Abbreviations: Hexose-P, hexose phosphates; Triose-P, triose phosphates; PEP, phospho*enol*pyruvate; OPPP, the oxidative pentose phosphate pathway; PEPC, phospho*enol*pyruvate carboxylase; PEPCK, phospho*enol*pyruvate carboxykinase; ME, malic enzyme; TCA cycle, the tricarboxylic acid cycle.

### Simultaneous operation of multiple metabolic processes involved in the cycling of metabolite storages over a diel cycle in guard cells

In the Open phase, the metabolic flux prediction for the standard scenario showed an import of K^+^ ions through the K^+^/H^+^ symporter and malate synthesis from multiple metabolic processes (Figure 2). The main carbon source for malate synthesis in this phase was the degradation of starch stored at night, contributing to 64% of the carbon accumulated in malate. The resulting hexose phosphate from starch degradation merges with the CBC, which is another carbon source for malate synthesis. The products of CBC were then converted into phospho*enol*pyruvate (PEP) through glycolysis. The third carbon source is the fixation of CO_2_ in the conversion of PEP into oxaloacetate (OAA) by PEP carboxylase (PEPC), contributing to 25% of the carbon in malate accumulation. OAA produced from PEP carboxylase was subsequently converted into malate by malate dehydrogenase.

The central role of PEP in malate synthesis predicted by our model corresponds to the findings in existing literature that PEP functions as the primary precursor to OAA. Santelia and Lawson (2016) summarised the numerous studies suggesting that PEPC catalyses the carboxylation of PEP to produce OAA and, subsequently, to produce malate (Willmer et al., 1973; Willmer and Dittrich, 1974; Outlaw and Kennedy, 1978; Rao and Anderson, 1983; Chollet et al., 1996). Robaina-Estevez et al. (2017) likewise predicted the use of PEPC in malate synthesis during stomata opening. Our findings showed that it is possible for carbon fixation from Rubisco to happen concurrently with PEPC phosphorylation, consistent with a stable isotope labelling study showing CO_2_ fixation via both Rubisco and PEPC in tobacco guard cells (Daloso et al., 2015).

In the Day phase, the model was constrained to take up sucrose (80% of the amount of sucrose required for maintaining the opening of the stomata) and export K^+^ from the guard cell. The model predicted the synthesis of sucrose and starch from malate degradation and carbon fixation from the Calvin-Benson cycle. All the accumulated malate from the Open phase was converted into PEP before undergoing gluconeogenesis to produce hexose phosphates for sucrose and starch synthesis. Flux variability analysis showed that NADP-malic enzyme (NADP-ME) and PEPCK were equally favourable (Data S2). This can explain the experimental evidences for malate decarboxylation by PEPCK (Penfield et al., 2012) and malic enzyme (Outlaw et al., 1981; Schnabl, 1981) in the guard cells of different plants. Though it is generally accepted that malate export is more important than metabolism for malate degradation within the guard cell (Dittrich and Raschke, 1977; Santelia and Lawson, 2016), our findings show that it is possible for the degradation of malate to contribute to sucrose and starch synthesis via gluconeogenesis. In addition to malate degradation, the CBC provided the rest of the carbon for sucrose and starch synthesis. This includes the refixation of CO_2_ produced from malate decarboxylation. While existing research overwhelmingly shows that guard cells prefer importing apoplastic sucrose produced by mesophyll photosynthesis for its various functions (Tarczynski et al., 1989; Reckmann et al., 1990; Vavasseur and Raghavendra, 2005; Santelia and Lawson, 2016), our model demonstrated a possible metabolic mode in which sucrose as osmolyte for keeping the stomata opened can be synthesised from stored malate and CBC.

In the Close phase, sucrose was degraded to form hexose phosphates, the majority of which was used for starch synthesis. The rest of the sucrose was metabolised through the oxidative part of the OPPP to produce NADPH for cellular maintenance and pentose phosphates. The pentose phosphates were then converted into hexose phosphates and triose phosphates through the non-oxidative part of OPPP in the plastid. The resulting hexose phosphates and triose phosphates were metabolised through glycolysis to produce pyruvate, which was transported into the mitochondrion and metabolised through pyruvate dehydrogenase and the TCA cycle, ultimately producing ATP by the mitochondrial electron transport chain for cellular maintenance. Existing research has suggested various non-osmoregulatory roles for sucrose, for instance, as a substrate for respiration or as a contributor of carbon skeletons for starch (Santelia and Lawson, 2016). Our findings support the view that sucrose can be used for both functions and suggest a viable metabolic route for each function.

In the Night phase, a large proportion (91%) of the starch accumulated from the Close phase was degraded to fuel the guard cell with ATP and NADPH for maintenance processes through glycolysis, the OPPP and the TCA cycle. The remaining starch (9%of starch stored during stomatal closure) was stored at the end of the Night phase and contributed to the production of malate during the Open phase as the stoma opens.

### Malate and Cl^-^ are similarly energetically efficient as counterion of K^+^

One of the unresolved questions about guard cell metabolism concern the choice of counterion for K^+^. To evaluate if the choice of either counterion is due to the respective energy costs to accumulate either counterion, we compared the energy costs (measured by photon influx) between the standard scenario (i.e. the “malate-only” scenario since malate was designated as the sole counterion) and the “Cl^-^-only” scenario where Cl^-^ was allowed to accumulate and malate accumulation was constrained to 0 with all other constraints being the same as the standard scenario. Interestingly, despite the fact that the malate-only scenario accumulates a larger amount of K^+^ than the Cl^-^-only scenario (0.75 vs. 0.5 mmol g^-1^ DW h^-1^ in the malate-only and Cl^-^-only scenarios respectively), we found that total photon influx was very similar (<0.2% difference) between the two scenarios (Table 1), suggesting that the energy requirement for importing K^+^ and Cl^-^ is similar to that of importing additional K^+^ with the counterion of malate being produced from starch degradation and photosynthesis.

**Table 1.**
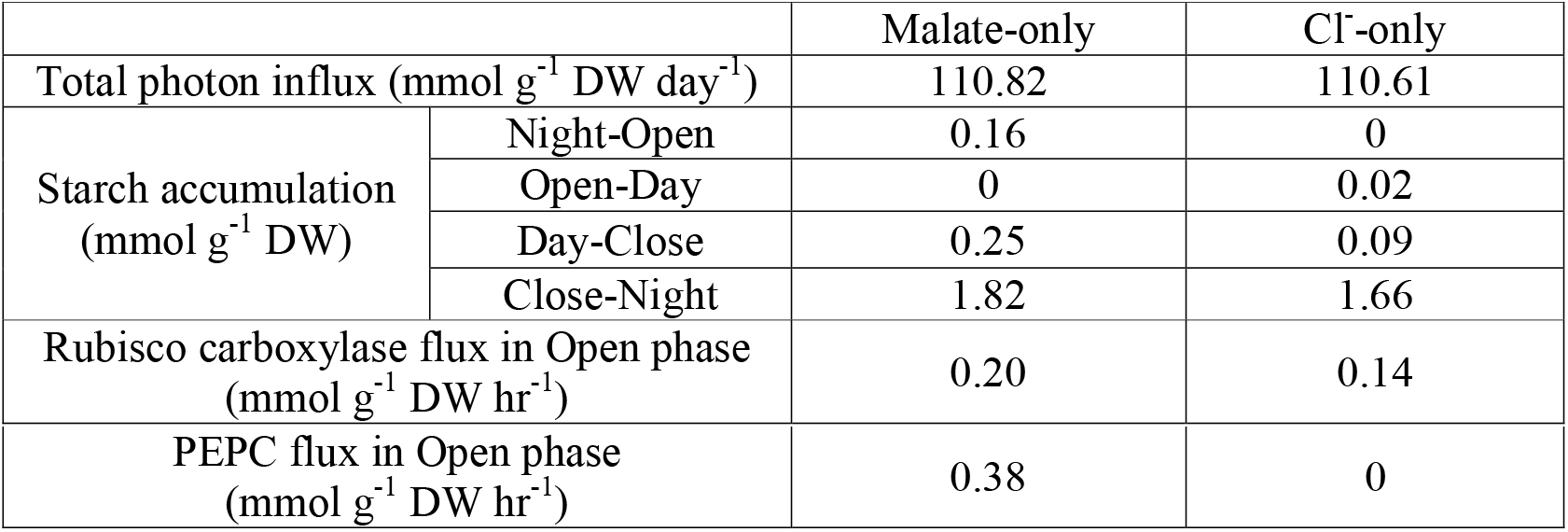
Comparison of energetics and fluxes in starch accumulation and carbon fixation between the malate-only and Cl^-^-only scenarios. Starch accumulation between two consecutive phases were denoted as X-Y where starch accumulates from phase X to phase Y.

Besides the obvious differences in the accumulation of Cl^-^, malate and K^+^, our model predicted two key differences between the flux modes of the malate-only and Cl^-^-only scenarios – i) timing of starch accumulation and ii) amount of carbon fixation during the Open phase (Table 1). In the malate-only scenario, starch from the Night phase is degraded to provide carbon for malate synthesis during the Open phase; whereas starch starts to accumulate during the Open phase in the Cl^-^-only scenario to provide carbon for the subsequent phases. In addition, the Cl^-^-only scenario accumulates less starch than the malate-only scenario in all other phases. This is a strong indication that starch accumulation contributes to the use of malate as an osmolyte for stomatal opening. Despite the absence of starch accumulation during the Open phase for the malate-only scenario, it was predicted to have a higher carbon fixation rate during the Open phase which is reflected in both higher Rubisco carboxylase flux and higher PEPC flux. This is in line with the previous results suggesting that both Rubisco and PEPC can contribute to malate synthesis for stomatal opening (Daloso et al., 2015).

### Flexibility of central carbon metabolism in guard cells with external sucrose import and photon influx

To investigate the effect of sucrose uptake from the apoplast on guard cell metabolism, we performed a scan of sucrose uptake (i.e. varying the magnitude of sucrose uptake flux) for the standard (malate-only) scenario (Data S3). Photon influx decreased as sucrose uptake increased (Figure 3a), which means sucrose could be used as complementary energy source to photon. Interestingly, there is a shift in gradient at sucrose uptake rate of 87 μmol g^-1^ DW h^-1^. At sucrose uptake rate below 87 μmol g^-1^ DW h^-1^, the Calvin-Benson cycle operates during the day to fix carbon for sucrose and starch (Figure 3b). As sucrose uptake rate increases above 87 μmol g^-1^ DW h^-1^, carbon fixation via the Calvin-Benson cycle stops during the day and the imported sucrose starts to be metabolised through the OPPP (Figure 3c). When sucrose uptake rate is above 175 μmol g^-1^ DW h^-1^, the imported sucrose alone is enough to drive guard cell metabolism for the whole diel cycle without any photon influx. This analysis demonstrated the flexibility of guard cell metabolism to varying photon influx and sucrose uptake by operating in a mixotrophic mode during the day with a switch from sucrose synthesis via the Calvin-Benson cycle to sucrose degradation via the OPPP.

**Figure 3.**
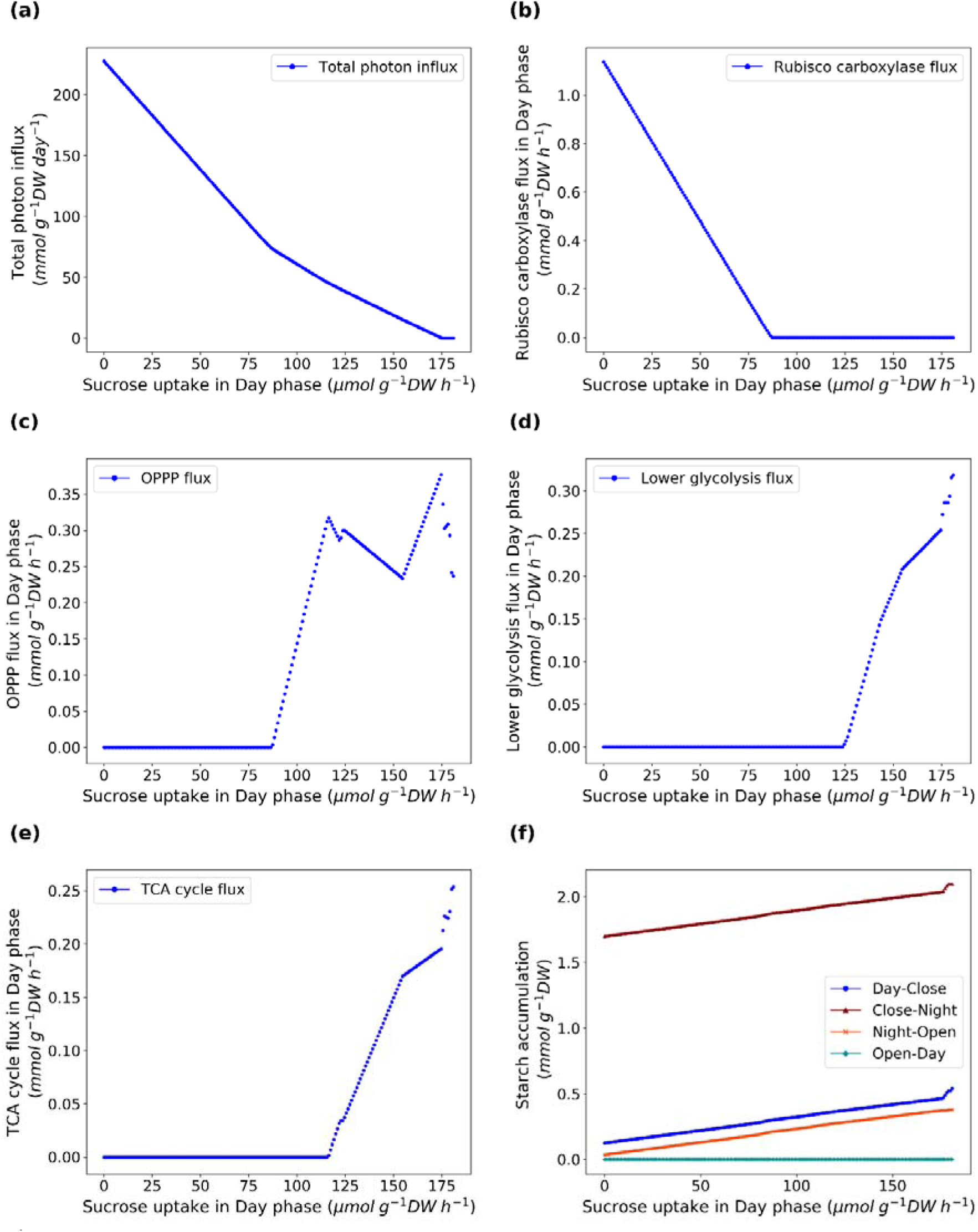
Changes in metabolic fluxes in response to sucrose uptake rate during the Day phase. (a) Total photon influx over a diel cycle; (b) Rubisco carboxylase flux during the Day phase; (c) Flux of the OPPP during the Day phase based on the sum of glucose-6-phosphate dehydrogenase fluxes in all compartments during the Day phase; (d) Flux of lower glycolysis during the Day phase based on the sum of pyruvate kinase fluxes in all compartments during the Day phase; (e) Flux of the TCA cycle during the Day phase based on the flux of succinate dehydrogenase during the Day phase; (f) Amount of starch accumulation from phase X to phase Y denotes as X-Y in the figure label.

At high sucrose uptake from the apoplast during the day, lower glycolysis and the TCA cycle becomes active during the day as the contribution of sucrose to the guard cell’s energy demand increases (Figure 3d,e). This is in-line with isotope labelling experiments of guard cells showing external sucrose being metabolised through glycolysis and the TCA cycle (Medeiros et al., 2018). Our analysis suggests that the role of sucrose in guard cell metabolism could vary depending on the availability and uptake of sucrose from the apoplast.

With our four-phase model, we were able to show that sucrose import during the Day phase can have impact on the metabolism of other phases. While the metabolic demand remained the same throughout the analysis, the model predicted an increase in starch accumulation from the Day phase to the Open phase through the Close and Night phases as sucrose uptake during the day increases (Figure 3f). This increases the contribution of starch degradation to malate synthesis during the Open phase as photon influx decreases. When the contribution of photon to malate synthesis during the Open phase is relatively high (i.e. low Day phase sucrose uptake), the model predicted an alternative flux mode of the Calvin-Benson cycle which involves diphosphate—fructose-6-phosphate 1-phosphotransferase (PFP) and transaldolase (Figure 4). The diversion of all dephosphorylation steps in the Calvin-Benson cycle to PFP allows the maximum amount of PPi production to drive the pumping of protons by the tonoplast H^+^-translocating pyrophosphatase. We predict that this mode of the Calvin-Benson cycle involving PFP and transaldolase could occur in cells that have a high PPi demand (e.g. pumping of metabolites across the tonoplast) and a low amount of alternative PPi sources (e.g. in non-growing cells that do not produce PPi from the polymerisations of proteins and nucleic acids). Besides guard cells, such conditions could potentially be found in other plant systems such as ripening fruits with photosynthetic activity. From the predicted flux mode, we hypothesise that an analogous flux mode with all dephosphorylation steps in the Calvin-Benson cycle diverted to sedoheptulose-1,7-bisphosphatase (Figure S1) which could operate depending on the regulation of the transaldolase and the dephosphorylating enzymes in the Calvin-Benson cycle.

**Figure 4.**
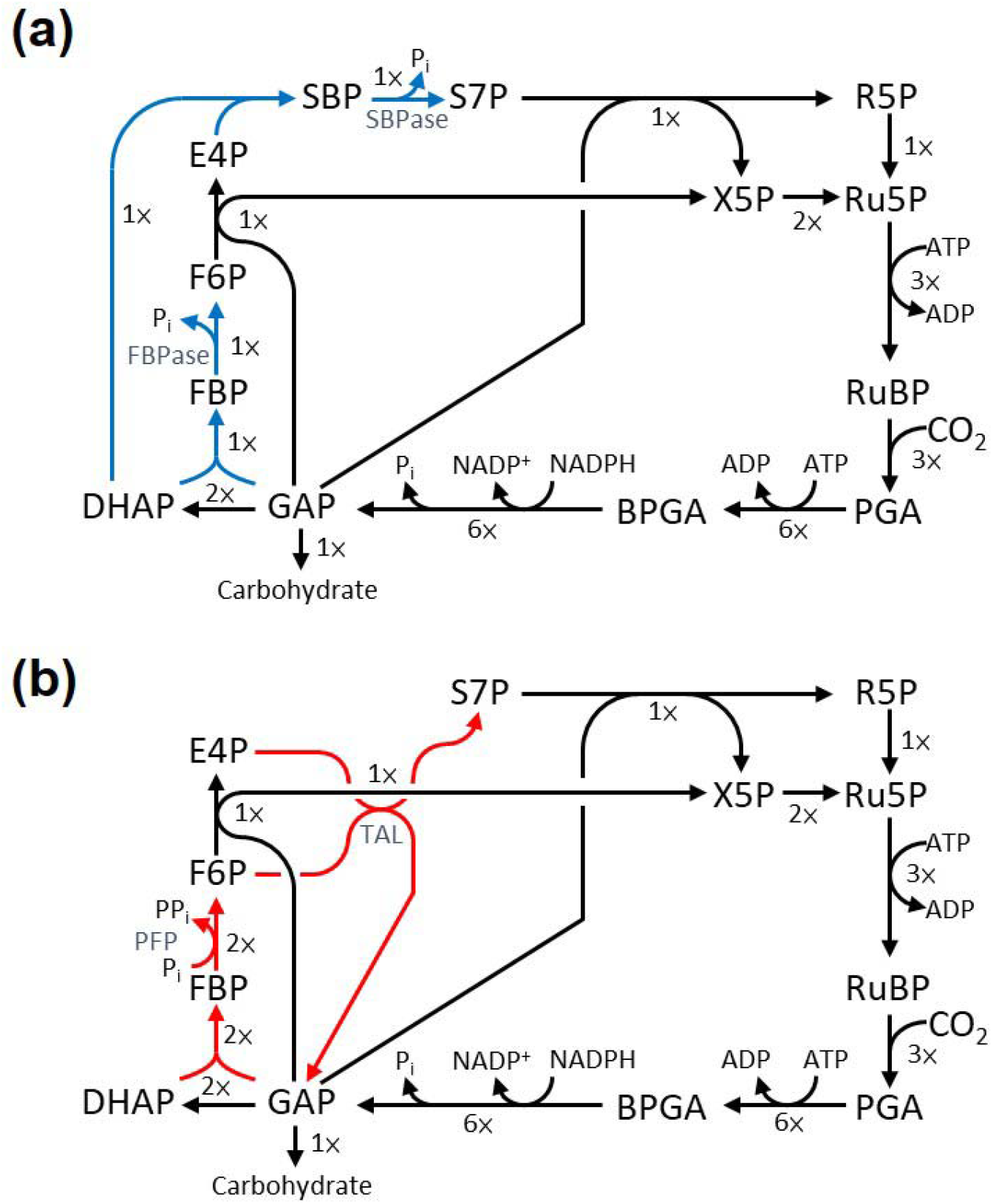
Alternative flux modes of the Calvin-Benson cycle with different dephosphorylating enzymes. (a) Traditional flux mode of the Calvin-Benson cycle with flux through both fructose-6-phosphatase (FBPase) and sedoheptulose-1,7-bisphosphatase (SBPase); (b) an alternative flux mode predicted during stomatal opening in guard cell with diphosphate—fructose-6-phosphate 1-phosphotransferase (PFP) as the sole dephosphorylating enzyme. Blue and red arrows represent reactions with different fluxes between the two flux modes. The number besides the arrow denotes the relative flux of the reaction. Metabolite abbreviations: PGA, 3-phosphoglycerate; BPGA, glycerate 1,3-bisphosphate; GAP, glyceraldehyde 3-phosphate; DHAP, dihydroxyacetone phosphate; FBP, fructose 1,6-bisphosphate; F6P, fructose 6-phosphate; E4P, erythrose 4-phosphate; SBP, sedoheptulose-1,7-bisphosphate; S7P, sedoheptulose 7-phosphate; R5P, ribose 5-phosphate; X5P, xylulose 5-phosphate; Ru5P, ribulose 5-phosphate; RuBP, ribulose 1,5-bisphosphate.

## Materials and methods

### Construction of a four-phase model for guard cell metabolism

A core plant metabolic model (Shameer et al., 2018) was converted into a single-phase model, which was then duplicated four times to model the four stages of guard cell behaviour - the opening and closing of stomata, and the intervening day and night phases (Appendix S1, S2). The names of reactions and metabolites of each of the four phases were labelled with phase-specific suffixes (“Open”, “Day”, “Close” and “Night”). Unidirectional storage reactions were added for K^+^, Cl^-^, malate, sucrose, starch and H^+^, where metabolites accumulated at the end of each phase were made available for consumption in the subsequent phase through storage reactions. Storage reactions were labelled with suffixes denoting the connected phases (e.g. “K_OpenDay_storage” for accumulation of K^+^ during the Open phase which can be consumed in the Day phase). The code for the construction of the four-phase guard cell model, including a list of minor modifications from the model in Shameer et al. (2018), can be found in Appendix S3. The model files and all scripts used in the study can be found in https://github.com/mauriceccy/guardcell/.

### Constraints for four-phase model of guard cell metabolism

Besides the constraints described in the Results section for modelling guard cell metabolism under different scenarios, the following constraints were applied for all simulations in this study. During light phases (Open and Day), photon influx was not constrained, whereas during the dark phases (Close and Night), photon influx was constrained to zero. The length of respective phases was modelled by constraining the ratio of total photon influx, with respect to a 1:11 Open to Day light phase, and by constraining the cellular maintenance costs in the ratio of 1:11:1:11 for Open, Day, Close and Night phases, assuming constant cellular maintenance cost per hour throughout the four phases. Cellular maintenance costs were modelled by constraining generic ATPase and NADPH oxidase reactions to carry a nonzero flux, in a 3:1 ATP to NADPH ratio, based on estimations from Arabidopsis heterotrophic cell culture under control conditions (Cheung et al., 2013). ATP consumption was set to take place only in the cytosol, whereas NADPH consumption was equally distributed between the cytosol, mitochondrion and plastid as in Cheung et al. (2013). The opening and closing of stoma were modelled by constraining the model to undergo a fixed change in osmotic pressure. This was done by setting a fixed sum of accumulation of osmolytes scaled by their osmotic coefficients. The value of 0.9 mosmol g^-1^ DW was used based on experimental measurements in *Vicia faba* guard cells (Rechmann et al., 1990). We applied osmolarity coefficients to each osmolyte within our modelling framework that correspond to the observed osmotic concentration of each osmolyte (the osmotic coefficients of KCl, K2malate and sucrose are 0.9, 0.8 and 1 respectively), such that the model would accumulate the appropriate amount of osmolytes to achieve the set osmotic pressure. To model photorespiration, Rubisco carboxylase to oxygenase ratio was constrained to a 3:1 ratio under normal atmospheric conditions (Gutteridge and Pierce, 2006). Chloroplastic NADPH dehydrogenase and plastoquinol oxidase were set to zero similar to previous studies (Cheung et al., 2014) given their negligible role in photosynthesis (Yamamoto et al., 2011; Josse et al., 2000). Only H_2_O, CO_2_, O_2_, K^+^ and Cl^-^ were allowed to freely enter and exit the cell. All other import, export and biomass reactions (except photon and sucrose as stated in their corresponding constraints) were set to carry zero flux.

### Flux balance analysis and flux variability analysis

Flux balance analysis was conducted using scobra (https://pypi.org/project/scobra/), an extension package of COBRApy (Ebrahim et al., 2013). In this study, minimising photon influx was used as the primary objective function. Minimisation of the absolute sum of fluxes was applied as a secondary set of constraints, per the assumption that guard cell metabolism would minimise its investment in enzyme machineries. Flux variability analysis was performed as described in Mahadevan and Schilling (2003). In brief, the objective value of the primary objective function was set as an additional constraint followed by the minimisation and maximisation of each reaction in the model. A flux range for each reaction was obtained and represented as (v_min_, v_max_), where v_min_ and v_max_ are the minimum and maximum flux values respectively.

## Acknowledgements

We thank T. C. R. Williams for discussion on guard cell biochemistry. This work was supported by Yale-NUS College (R-607-265-233-121). The authors declare they have no conflict of interest.

## Supporting Information

**Appendix S1.** Four-phase model of guard cell metabolism in SBML format.

**Appendix S2.** Four-phase model of guard cell metabolism in Excel format.

**Appendix S3.** Instruction and code for constructing the four-phase guard cell model.

**Data S1.** Flux solutions of the individual simulated scenarios from the four-phase guard cell model using parsimonious flux balance analysis.

**Data S2.** Flux range of the individual simulated scenarios from the four-phase guard cell model using flux variability analysis.

**Data S3.** Flux solutions of sucrose uptake scan from the four-phase guard cell model using parsimonious flux balance analysis.

**Figure S1.** Alternative flux modes of the Calvin-Benson cycle with different dephosphorylating enzymes.

