## Appendix S3 for "A multi-phase flux balance model reveals flexibility of guard cell central carbon metabolism"

**Appendix S2**

Instruction and code for constructing the four-phase guard cell model

Besides the details listed in the main text of the manuscript, a few minor corrections/modifications were applied to the core plant metabolic model in Shameer et al., (2018).

- The reaction “HEXOKINASE_RXN_MANNOSE_c” was corrected to have all its metabolites in the cytosol with stoichiometry of the various protonated states correct to the cytosolic values.
- The constraint for a cytosolic proton leak reaction “unlProtHYPO_c” was set to 0.
- Proton transport between the cytosol and the mitochondrial intermembrane space “H_ic” was set to reversible. It was tested to have negligible effect on the flux solutions in this study.
- Irreversible proton transport reactions were added between the cytosol and the vacuole (“H_vc”) and between the extracellular space and the cytosol (“H_ec”).
- Transport reactions for K^+^, Cl^-^ and malate across the plasma membrane and the tonoplast were added (“K_rev_ec”, “K_vc”, “Cl_tx”, “Cl_ec”, “Cl_rev_ec”, “Cl_vc”, “Cl_rev_vc”, “MAL_tx”, “MAL_ec”, “MAL_rev_ec”) to the model.
- Storage reactions for K^+^, malate, starch, sucrose, Cl^-^ and H^+^ were added to the model.

The code below required scobra (<https://pypi.org/project/scobra/>) where "CoreLeaf.xml" is the core plant metabolic model from Shameer et al. (2018) and “GC.xls” and “GC.xml” are the four-phase guard cell model in Excel and SBML formats.

```
import scobra
m = scobra.Model("CoreLeaf.xml")

# Make a single model
for r in m.Reactions():
    if r.startswith("EX_"):
        m.DelReaction(r)
    elif r.endswith("2"):
        m.DelReaction(r)
    elif r.endswith("dielTransfer"):
        m.DelReaction(r)
m.DelReaction("diel_biomass")

for met in m.Metabolites():
    if met.endswith("_boundary"):
        m.DelMetabolite(met)
    elif met.endswith("2"):
        m.DelMetabolite(met)
    elif met.endswith("_t"):
        m.DelMetabolite(met)

new_model = m.copy()
for reac in new_model.reactions:
    reac.id = reac.id[:-1]
for met in new_model.metabolites:
    met.id = met.id[:-1]
new_model.repair()
m = new_model

for r in m.reactions:
    if "SUBSYSTEM" in r.notes:
        r.subsystem = r.notes["SUBSYSTEM"]
    r.notes = {}

# Fix HEXOKINASE_RXN_MANNOSE_c
m.DelReactions(["HEXOKINASE_RXN_MANNOSE_c"])
m.AddReaction("HEXOKINASE_RXN_MANNOSE_c", {"ATP_c":-0.65, "MANNOSE_c":-1, "aATP_c":-0.35, "ADP_c":0.5, "MANNOSE_6P_c":1, "PROTON_c":0.85, "aADP_c":0.5}, rev=False)

# Set unlProtHYPO to zero
m.SetConstraints({'unlProtHYPO_c':(0,0)})

# Set H_ic to be reversible
m.SetConstraints({'H_ic':(None,None)})

# Add irreversible H_vc
m.AddReaction("H_vc",{"PROTON_v":-1, "PROTON_c":1}, rev=False)

# Add irreversible H_ec
m.AddReaction("H_ec",{"PROTON_e":-1, "PROTON_c":1}, rev=False)

# Add K+ reactions
m.AddReaction('K_rev_ec', {'KI_c':-1, 'KI_e':1}, rev = False)
m.AddReaction('K_vc', {"KI_v":-1, "KI_c":1}, rev = False)

# Add Cl reactions
m.AddReaction('Cl_tx', {'CL_e' : 1}, rev = True)
m.AddReaction('Cl_ec', {'PROTON_e':-2, 'CL_e':-1, 'PROTON_c':2, 'CL_c':1}, rev = False)
m.AddReaction('Cl_rev_ec', {'CL_c':-1, 'CL_e':1}, rev = False)
m.AddReaction('Cl_vc', {'CL_v':-1, 'CL_c':1}, rev = False)
m.AddReaction('Cl_rev_vc', {'CL_c':-2, 'PROTON_v':-1, 'CL_v':2, 'PROTON_c':1}, rev = False)

# Add Malate reactions
m.AddReaction('MAL_tx', {'MAL_e' : 1}, rev = True)
m.AddReaction('MAL_ec', {'PROTON_e':-3, 'MAL_e':-1, 'PROTON_c':3, 'MAL_c':1}, rev = False)
m.AddReaction('MAL_rev_ec', {'MAL_c':-1, 'MAL_e':1}, rev = False)

big_m = m.DuplicateModel({"_Open", "_Day", "_Close", "_Night"}) #suffixes is a dir of strings

##Storage Reactions
#for K
big_m.AddReaction('K_OpenDay_storage', {'KI_v_Open' : -1, 'KI_v_Day': 1}, rev = False)
big_m.AddReaction('K_DayClose_storage', {'KI_v_Day' : -1, 'KI_v_Close': 1}, rev = False)
big_m.AddReaction('K_CloseNight_storage', {'KI_v_Close' : -1, 'KI_v_Night': 1}, rev = False)
big_m.AddReaction('K_NightOpen_storage', {'KI_v_Night' : -1, 'KI_v_Open': 1}, rev = False)

#for malate
big_m.AddReaction('Malate_OpenDay_storage', {'MAL_v_Open': -0.7, 'aMAL_v_Open': -0.3, 'MAL_v_Day': 0.7, 'aMAL_v_Day': 0.3}, rev = False)
big_m.AddReaction('Malate_DayClose_storage', {'MAL_v_Day': -0.7, 'aMAL_v_Day': -0.3, 'MAL_v_Close': 0.7, 'aMAL_v_Close': 0.3}, rev = False)
big_m.AddReaction('Malate_CloseNight_storage', {'MAL_v_Close': -0.7, 'aMAL_v_Close': -0.3, 'MAL_v_Night': 0.7, 'aMAL_v_Night': 0.3}, rev = False)
big_m.AddReaction('Malate_NightOpen_storage', {'MAL_v_Night': -0.7, 'aMAL_v_Night': -0.3, 'MAL_v_Open': 0.7, 'aMAL_v_Open': 0.3}, rev = False)

#for starch
big_m.AddReaction('Starch_OpenDay_storage', {'STARCH_p_Open': -1, 'STARCH_p_Day': 1}, rev = False)
big_m.AddReaction('Starch_DayClose_storage', {'STARCH_p_Day': -1, 'STARCH_p_Close': 1}, rev = False)
big_m.AddReaction('Starch_CloseNight_storage', {'STARCH_p_Close': -1, 'STARCH_p_Night': 1}, rev = False)
big_m.AddReaction('Starch_NightOpen_storage', {'STARCH_p_Night': -1, 'STARCH_p_Open': 1}, rev = False)

#for sucrose
big_m.AddReaction('Sucrose_OpenDay_storage', {'SUCROSE_v_Open': -1, 'SUCROSE_v_Day': 1}, rev = False)
big_m.AddReaction('Sucrose_DayClose_storage', {'SUCROSE_v_Day': -1, 'SUCROSE_v_Close': 1}, rev = False)
big_m.AddReaction('Sucrose_CloseNight_storage', {'SUCROSE_v_Close': -1, 'SUCROSE_v_Night': 1}, rev = False)
big_m.AddReaction('Sucrose_NightOpen_storage', {'SUCROSE_v_Night': -1, 'SUCROSE_v_Open': 1}, rev = False)

#for cl
big_m.AddReaction('Cl_OpenDay_storage', {'CL_v_Open': -1, 'CL_v_Day': 1}, rev = False)
big_m.AddReaction('Cl_DayClose_storage', {'CL_v_Day': -1, 'CL_v_Close': 1}, rev = False)
big_m.AddReaction('Cl_CloseNight_storage', {'CL_v_Close': -1, 'CL_v_Night': 1}, rev = False)
big_m.AddReaction('Cl_NightOpen_storage', {'CL_v_Night': -1, 'CL_v_Open': 1}, rev = False)

# Add Proton_v_dielTransfer
big_m.AddReaction('H_OpenDay_storage', {'PROTON_v_Open': -1, 'PROTON_v_Day': 1}, rev = False)
big_m.AddReaction('H_DayClose_storage', {'PROTON_v_Day': -1, 'PROTON_v_Close': 1}, rev = False)
big_m.AddReaction('H_CloseNight_storage', {'PROTON_v_Close': -1, 'PROTON_v_Night': 1}, rev = False)
big_m.AddReaction('H_NightOpen_storage', {'PROTON_v_Night': -1, 'PROTON_v_Open': 1}, rev = False)

####How to write model####
big_m.WriteModel("GC.xls")
big_m.WriteModel("GC.xml")
```
