## Supplementary material for "A multi-phase flux balance model reveals flexibility of guard cell central carbon metabolism": Figure S1

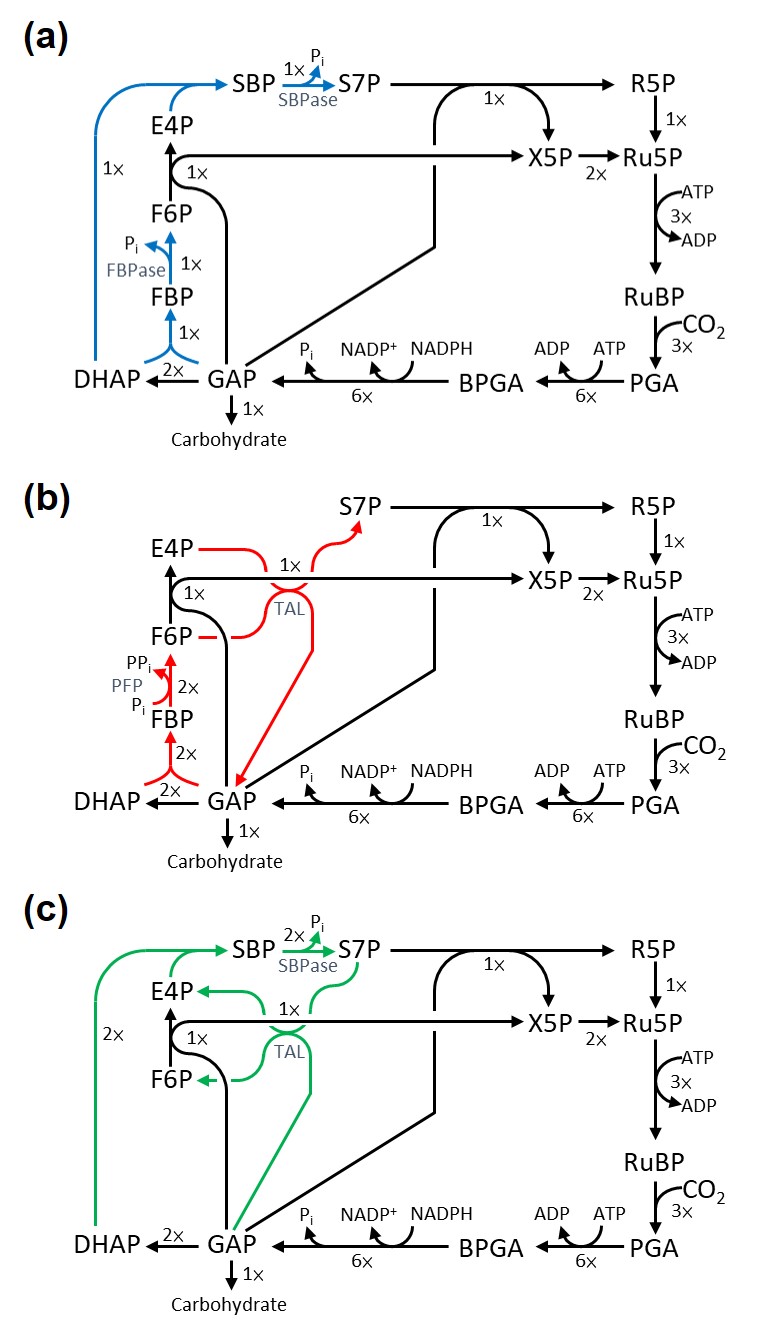

**Figure S1. Alternative flux modes of the Calvin-Benson cycle with different dephosphorylating enzymes.** (a) Traditional flux mode of the Calvin-Benson cycle with flux through both fructose-6-phosphatase (FBPase) and sedoheptulose-1,7-bisphosphatase (SBPase); (b) an alternative flux mode predicted during stomatal opening in guard cell with diphosphate—fructose-6-phosphate 1-phosphotransferase (PFP) as the sole dephosphorylating enzyme; (c) a hypothetical flux mode with SBPase as the sole dephosphorylating enzyme. Blue, red and green arrows represent reactions with different fluxes between the three flux modes. The number besides the arrow denotes the relative flux of the reaction. Metabolite abbreviations: PGA, 3-phosphoglycerate; BPGA, glycerate 1,3-bisphosphate; GAP, glyceraldehyde 3-phosphate; DHAP, dihydroxyacetone phosphate; FBP, fructose 1,6-bisphosphate; F6P, fructose 6-phosphate; E4P, erythrose 4-phosphate; SBP, sedoheptulose-1,7-bisphosphate; S7P, sedoheptulose 7-phosphate; R5P, ribose 5-phosphate; X5P, xylulose 5-phosphate; Ru5P, ribulose 5-phosphate; RuBP, ribulose 1,5-bisphosphate; G6P, glucose 6-phosphate.
